# Radiotherapy induces persistent innate immune reprogramming of microglia into a primed state

**DOI:** 10.1101/2023.05.10.539581

**Authors:** Daniëlle C. Voshart, Takuya Oshima, Yuting Jiang, Gideon P. van der Linden, Anna P. Ainslie, Luiza Reali Nazario, Fleur van Buuren-Broek, Ayla C. Scholma, Nieske Brouwer, Jeffrey Sewdihal, Uilke Brouwer, Rob P. Coppes, Inge R. Holtman, Bart J.L. Eggen, Susanne M. Kooistra, Lara Barazzuol

## Abstract

More than half of all brain tumour survivors experience debilitating and often progressive cognitive decline after treatment with radiotherapy. Microglia, the resident macrophages in the brain, have been implicated in this decline. In response to various insults microglia can develop innate immune memory (IIM), which can either enhance (priming) or repress (tolerance) the response to subsequent inflammatory challenges. Here, we investigated whether radiation affects the IIM of microglia by irradiating the brains of rats and later exposing them to a secondary inflammatory stimulus. Comparative transcriptomic profiling and protein validation of microglia isolated from irradiated rats showed a stronger immune response to a secondary inflammatory insult demonstrating that radiation can lead to long-lasting molecular reprogramming of microglia. Transcriptomic analysis of post-mortem normal-appearing non-tumour brain tissue of glioblastoma patients indicates that radiation-induced microglial priming is conserved in humans. Targeting microglial priming after radiotherapy or avoiding further inflammatory insults could decrease radiotherapy-induced neurotoxicity.

## Introduction

The number of long-term brain tumour survivors is steadily growing. However, 50 to 90% of surviving patients experience an irreversible and often progressive impairment in neurocognitive function at a detrimental cost to quality of life^1^. The associated neurocognitive sequelae include a decline in different domains, such as memory, attention, executive function, and intellectual abilities, particularly in survivors diagnosed at a younger age^2,3^. The degree and pattern of cognitive side effects vary among patients. Although tumour characteristics, age, surgical resection, and chemotherapy are known contributing factors, cognitive impairment has been mostly associated with radiotherapy treatment^2–4^. Radiotherapy is a life-saving treatment in the management of primary brain tumours. However, the normal healthy brain tissue is also exposed to radiation when the target tumour volume is irradiated, leading to short and long-term side effects^5^.

Microglia, the resident macrophages of the central nervous system, have been previously shown to play a role in the development of radiation-induced neurocognitive dysfunction^6–9^. Indeed, studies have reported that radiation can affect both the number and activity of microglia^10,11^. However, the long-term molecular and functional state of microglia after brain irradiation remains largely unknown. Similarly to other macrophages, microglia have been recently shown to develop innate immune memory (IIM) capacity^12,13^. After an initial insult IIM comprises long-lasting molecular reprogramming that translates into a functional state in which microglia either enhance (priming) or suppress (tolerance) their response to a subsequent inflammatory challenge^14,15^. The concept of microglial priming has been associated with ageing and many neurodegenerative diseases, and the resultant excessive immune response linked to neuropathology and cognitive decline^12,16,17^. Primed microglia were first demonstrated in a study showing exaggerated proinflammatory cytokine expression in a model of murine prion disease after a systemic endotoxin lipopolysaccharide (LPS) immune challenge^18^ and have been observed in several disease mouse models, including Alzheimer’s disease^19^ and accelerated ageing with defective DNA repair^14,20^. The transcriptomic profile of primed microglia was characterised by high expression of genes associated with phagosome, lysosome, and antigen presentation^21^, similar to the disease-associated microglia (DAM) profile^22^. In contrast to priming, microglia can also become desensitised to a subsequent immune insult entering an IIM state of tolerance, for instance after repeated doses of LPS^12,13^. Whether radiation can affect the IIM of microglia by leading to either immune priming or tolerance upon a secondary immune challenge has not been addressed before.

In this study, we showed by transcriptional profiling that radiation induces microglial priming resulting in an enhanced inflammatory response to LPS. Additionally, we performed a temporal and dose-dependent analysis and identified that radiation exposure causes persistent expression of microglial priming genes. The extent of this response is not affected by the administration of multiple smaller radiation doses. Importantly, a comparative transcriptomic analysis with post-mortem normal-appearing non-tumour brain regions of patients with glioblastoma (GBM) that were treated with radiotherapy revealed an overlap in radiation-associated microglial priming genes, indicating that this response is conserved in humans.

## Results

### Brain irradiation leads to microglial priming

To investigate the effect of radiation on the IIM of microglia, we irradiated the entire rat brain with 14 Gy X-rays and after 6 weeks exposed them to PBS vehicle or an endotoxin lipopolysaccharide (LPS) immune challenge. After 4 hours, CD11b^pos^/CD45^int^ microglia were isolated by fluorescence-activated cell sorting (FACS) and RNA sequencing was performed (RNA-seq) (Figure 1A, S1A). Principal component analysis (PCA) showed segregation of all four groups and distinct transcriptomic profiles (Figure 1B). We performed differentially expressed gene (DEG) analysis between the groups (Figure 1C, Figure S1B, Table S1). All the DEGs derived from the 4 comparisons were grouped into five different clusters by Manhattan distance-based hierarchical clustering (Figure 1D, S1C, Table S2). The largest number of genes was in cluster 4, which consisted of genes upregulated in the irradiated groups. Interestingly, cluster 4 contained priming signature genes, such as *Itgax* and *Gpnmb* (Figure 1D). Gene ontology (GO) analysis showed an enrichment of terms related to inflammation and immune regulation (Figure 1E). Whereas, cluster 5, consisting of genes downregulated after irradiation, was enriched for genes related to cell morphology and extracellular matrix organisation, as well as marker genes of homeostatic microglia that are known to be downregulated in reactive or primed microglia, including *Tmem119* (Figure 1D, 1E). Together, clusters 4 and 5 indicate that radiation induces transcriptional changes consistent with microglial priming.

**Figure 1:**
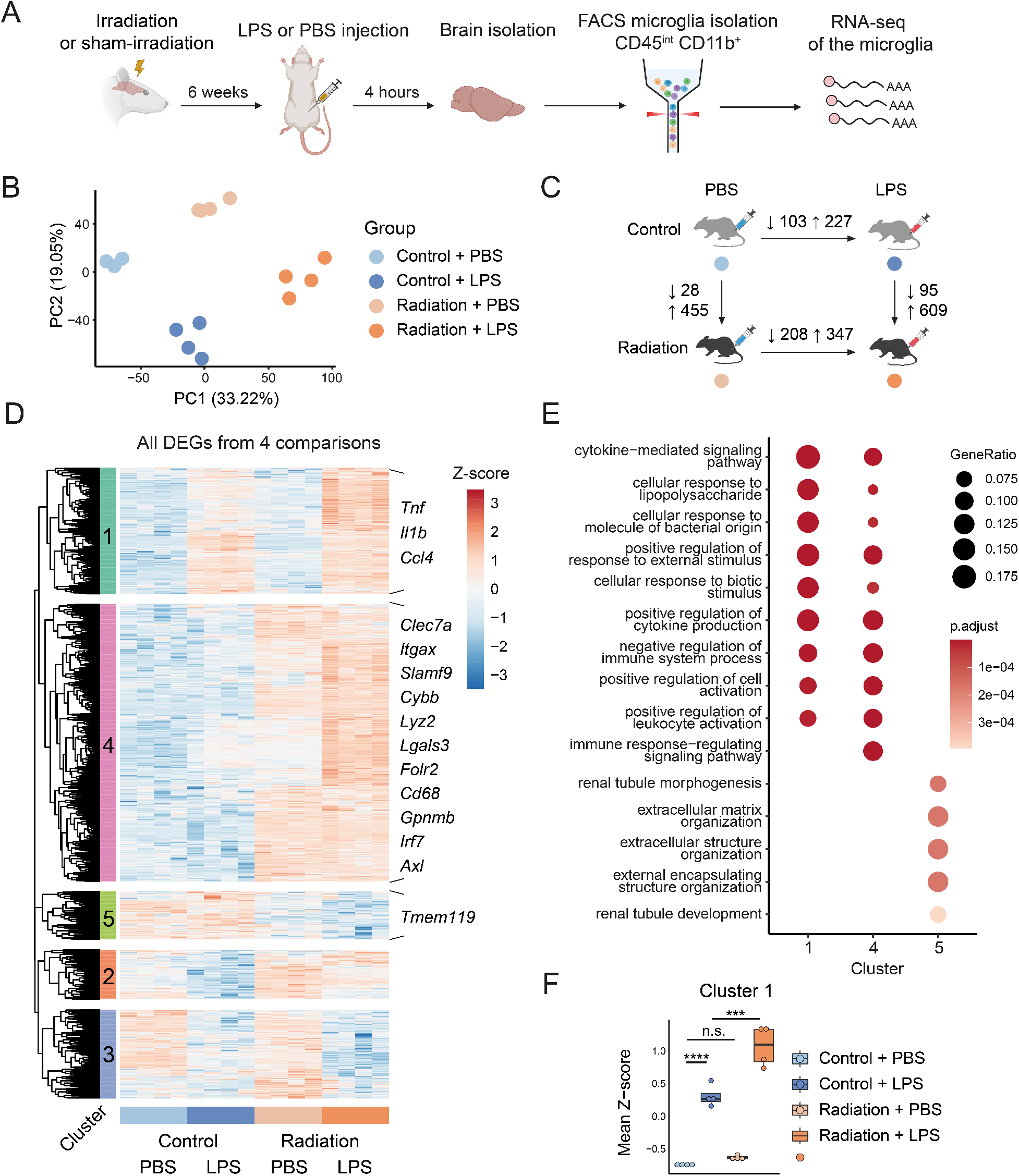
RNA-seq of microglia from irradiated rats subsequently challenged with LPS treatment. (A) Schematic overview of the experiment. Rats were (sham-)irradiated and subsequently challenged with PBS or LPS. Microglia were isolated by FACS after tissue dissociation of the whole brain and myelin removal, and then analyzed by RNA-seq. (B) Principal component analysis (PCA) of gene expression profiles of microglia in each condition (*n* = 4 per group). (C) The numbers of differentially expressed genes (DEGs) (Log_2_ fold change (LFC) > 2, adjusted p-value < 0.05) in the comparisons between the conditions. (D) Heatmap depicting gene expression of all the DEGs detected in the 4 comparisons in figure 1C. Hierarchical clustering resulted in 5 clusters of genes based on the expression pattern. Colours indicate row Z-score on the normalized read counts across samples for each gene. (E) Dot plot depicting Gene Ontology (GO) enrichment analysis (TOP 5 enriched functions of biological processes (BP)) of the genes in clusters 1, 4, and 5. (F) Box plots depicting the mean Z-score of genes in the clusters in each condition. Boxes are from the first quartile to the third quartile, and lines indicate the median. *n* = 4 animals per group. *** p <0.001, **** p <0.0001. Two-way ANOVA followed by Tukey’s multiple comparisons test was used for the comparison of the mean differences of multiple groups.

We next examined whether irradiated microglia react with a stronger response to a subsequent inflammatory challenge with LPS, which would confirm that microglia acquired a primed state after irradiation. Cluster 1 genes, including proinflammatory genes, such as *Tnf, Il1b*, and *Ccl4* were upregulated after LPS treatment and were significantly more upregulated in microglia derived from animals previously irradiated and exposed to LPS treatment compared to non-irradiated LPS-treated animals (Figure 1D, 1F). GO analysis demonstrated enrichment of biological processes related to inflammation and the general immune response (Figure 1E).

To further corroborate our findings, we compared our data to previously published microglial gene expression signatures associated with an acute, general, and primed response^21^ (Figure 2A). Higher expression of acute response genes was observed in microglia isolated from LPS-treated animals, while microglia isolated from irradiated animals show a higher expression of genes associated with the primed and mainly primed gene hubs described in Holtman et al.^23^ Microglial priming was further validated at the protein level by immunofluorescent staining of priming gene *Gpnmb*. At six weeks post irradiation, GPNMB was significantly more expressed in microglia of irradiated rats compared to controls (Figure 2B, 2C). Altogether, we revealed that brain irradiation induces microglial priming resulting in amplified immune responses upon secondary inflammatory insults.

**Figure 2:**
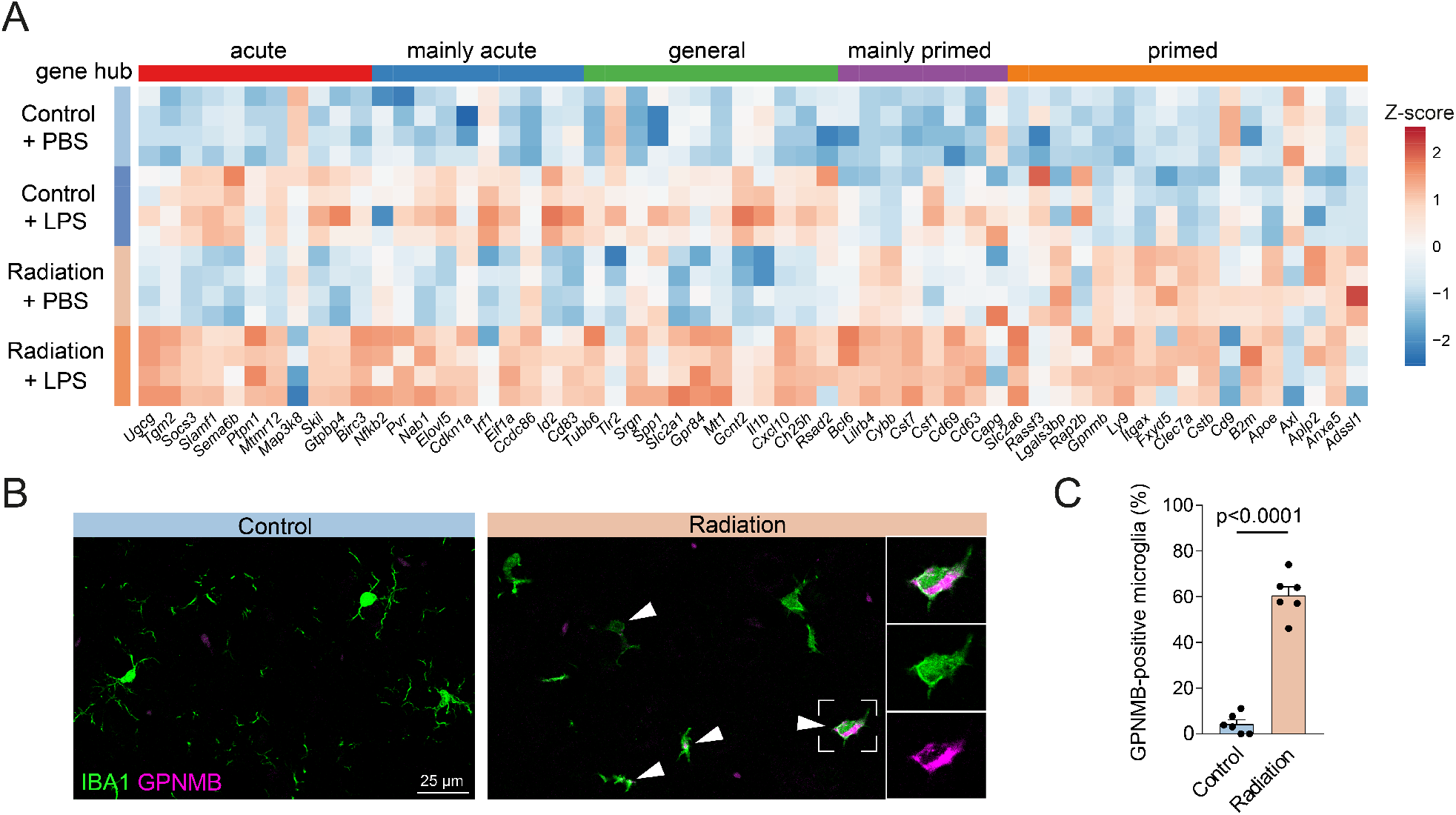
Transcriptional and protein validation of radiation-associated microglial priming. (A) Heatmap depicting expression of the genes associated with the acute immune response and the microglial priming described in literature^23^. Colours indicate column Z-score on the normalized read counts across samples for each gene. (B) Representative images of GPNMB (magenta) in microglia (IBA1, green), and (C) quantification of the percentage of GPNMB-positive microglia in the cortex of control and irradiated rats at 6 weeks post irradiation. Bar graph shows mean ±SEM. A two-tailed T-test was performed to compare the two groups. *n* = 6 animals per group.

### Brain irradiation induces a persistent microglial priming gene signature

Cognitive impairment in brain tumour survivors is considered to be irreversible and progressive in nature^1,24^. To investigate the persistence of radiation-induced microglial priming and whether this response could contribute to the late radiation effects, we quantified the expression of several microglial priming genes in rat cortical tissue at different time points post irradiation by qPCR (Figure 3A). We observed increased expression of most priming genes up to 50 weeks post irradiation, indicating that radiation leads to long-lasting molecular reprogramming of microglia (Figure 3A, S2A).

**Figure 3:**
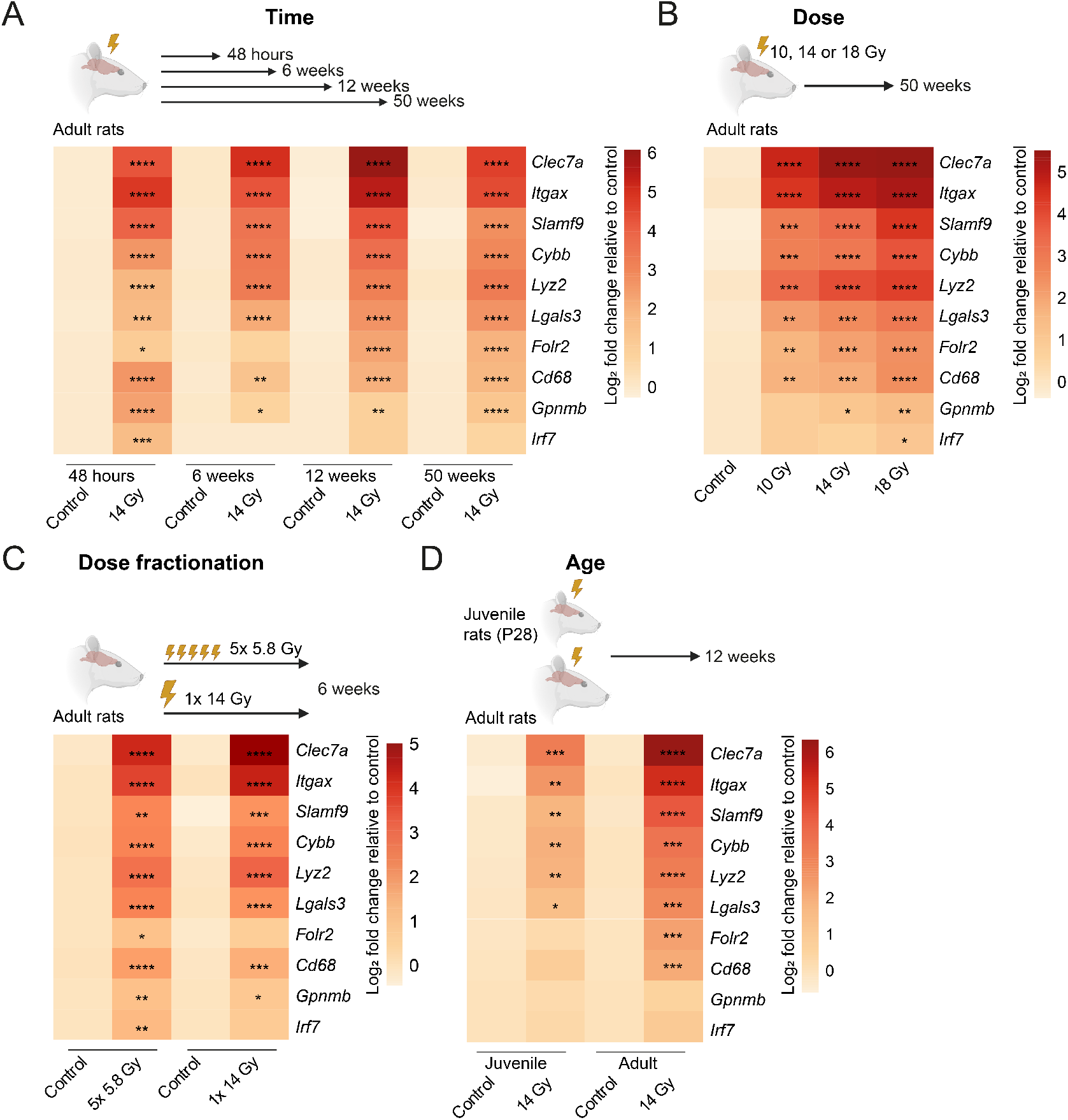
Persistence and dose-response effect of radiation on microglial priming genes. (A) Heatmap depicting gene expression of selected microglial priming genes of control rats and rats irradiated with 14 Gy at different time points after irradiation (48 hours, 6 week, 12 weeks, 50 weeks). (B) Heatmap depicting gene expression of selected microglial priming genes of control rats and rats irradiated with different radiation doses (10 Gy, 14 Gy, 18 Gy) at 50 weeks post irradiation. (C) Heatmap depicting gene expression of selected microglial priming genes of control rats and rats irradiated with a single radiation dose of 14 Gy or 5 smaller daily doses of 5.8 Gy at 6 weeks post irradiation. (D) Heatmap depicting gene expression of selected microglial priming genes of rats at two different age stages at the time of irradiation (P28 (4-week-old) juvenile, 9-week-old adult). Colours indicate log_2_ transformed mean of fold change relative to corresponding controls. **** p <0.0001, *** p <0.001, ** p <0.01, * p <0.05. Two-way ANOVA followed by Sidak’s multiple comparisons test with log_2_ transformed data (*n* = 4 animals per group) (A), one-way ANOVA followed by Tukey’s multiple comparisons test with log_2_ transformed data (*n* = 5-6 animals per group) (B), two-way ANOVA followed by Tukey’s multiple comparisons test with log_2_ transformed data (*n* = 5-6 animals per group) (C, D) were performed. See also Figure S2 and S3 for further details and statistics.

Next, to examine whether radiation-induced microglial priming also occurs to the same extent at different radiation doses, we quantified the expression level of the same microglial priming genes in rat cortical tissue at 50 weeks post-irradiation with 10, 14, and 18 Gy (Figure 3B). Most microglial priming genes were significantly upregulated in the 10 Gy irradiated animals, indicating that a relatively lower radiation dose is sufficient to induce microglial priming. The expression levels of these genes seem to increase with the radiation dose, with *Gpnmb* and *Irf7* showing a statistically significant upregulation in rats irradiated with a higher dose (Figure 3B, S2B).

We then investigated the effect of radiation dose fractionation on microglial priming gene expression, as radiotherapy treatment in brain tumour patients is normally delivered in multiple lower radiation doses. We irradiated rats with 5 daily fractions of 5.8 Gy and compared them to an equivalent single fraction dose of 14 Gy (Figure 3C). Overall, these two treatment schedules led to a similar increase in priming gene expression (Figure 3C, S3A).

Altogether these data provide evidence the radiation leads to persistent microglial priming and that its extent seems to be dose dependent and not altered by radiation dose fractionation.

### Juvenile rats show a less pronounced microglial priming response

Survivors of paediatric brain tumours are at a high risk of developing radiotherapy-induced cognitive decline^2^. To investigate whether microglial priming can also occur in juvenile rats, we irradiated the whole brain of postnatal day 28 (P28) rats with 14 Gy and assessed the expression of microglial priming genes in cortical tissue at 12 weeks post irradiation (Figure 3D). While most priming genes were significantly upregulated in both juvenile and adult rats compared to control, the relative expression of *Clec7a, Lyz2, Slamf9*, and *Folr2* was significantly higher in adult rats compared to juvenile rats (Figure 3D, S3B). This suggests that in juvenile rats the effect of radiation on microglial priming is less pronounced.

### Microglial priming occurs in the brain of GBM patients treated with radiotherapy

To identify the effect of radiotherapy on microglia in the human brain, we analysed RNA-seq data of normal appearing brain tissue from GBM patients that were treated with radiotherapy (NA-GBM, Table S4)^25^. Gene set enrichment analysis (GSEA) of DEGs enriched in NA-GBM showed a strong association with terms related to immune responses and cytokine production, indicating the contribution of reactive microglia in GBM patients treated with radiotherapy (Figure 4A). This was likely not an acute microglial response, as the cluster 1 genes, associated with an acute microglial LPS response (Figure 1D), were not increased in NA-GBM samples (Figure S4). In contrast, cluster 4 genes, associated with a radiation-induced response and encompassing microglial priming signature genes (Figure 1D), were more abundantly expressed in NA-GBM bulk tissue (Figure 4B). To confirm the transcriptional data at the protein level, we performed immunofluorescent staining for GPNMB in control and NA-GBM brain samples. Colocalization of IBA1 and GPNMB was significantly increased in NA-GBM brain samples (Figure 4C, 4D). These data suggest that microglial priming might occur in the brain of GBM patients in response to radiotherapy treatment.

**Figure 4:**
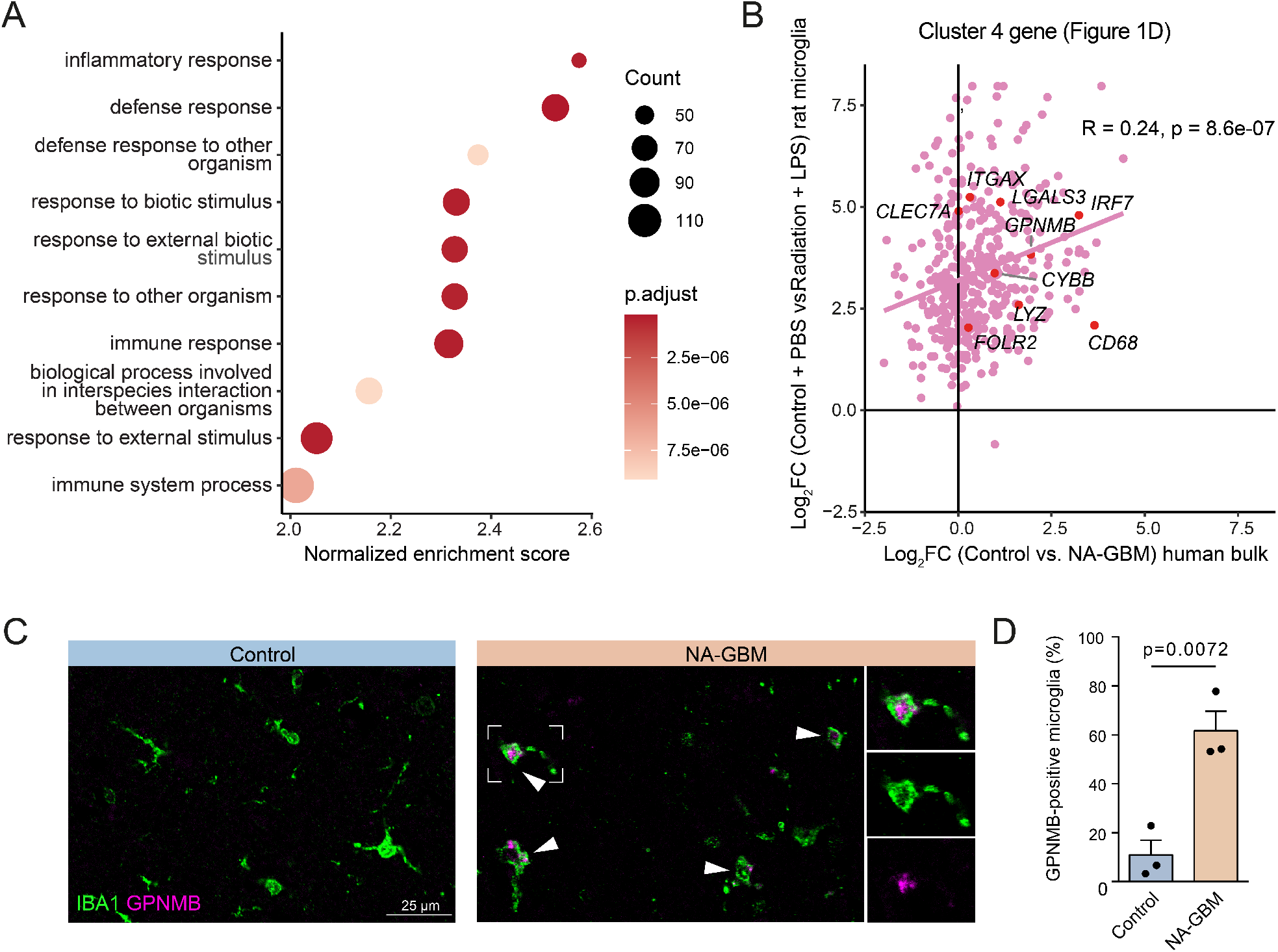
Microglial priming in GBM patients treated with radiotherapy. (A) GSEA (Biological Process) dot plot of DEGs (FC > 1.5, FDR < 0.05, RNAseq data derived from Ainslie et al.^25^) enriched in GBM patients treated with radiotherapy (NA-GBM) compared to neurotypical control individuals. (B) Scatter plot depicting a correlation of gene expression enrichment of cluster 4 genes in Figure 1D between the rat microglia-specific transcriptomic dataset (Control + PBS vs. Radiation + LPS) and the human bulk transcriptomic dataset (Control vs. NA-GBM). F-statistics was used to test the significance of regression coefficients in the linear regression model. (C) Representative images of GPNMB protein expression in microglia (IBA1) and (D) quantification of the percentage of GPNMB-positive microglia in control and NA-GBM post-mortem brain tissues. Bar graph shows mean ±SEM. A two-tailed T-test was performed to compare the two groups. *n* = 3 individuals per group. F-statistics was used to test the significance of regression coefficients in linear regression models.

## Discussion

Although radiotherapy is a life-saving treatment for both paediatric and adult brain tumour patients, it is associated with a high risk of developing long-term side effects. Microglia have been shown to play a role in radiotherapy-induced cognitive decline. However, the effect of radiation on the IIM and the resulting molecular state of microglia remain largely unknown. Here, by performing microglial transcriptional analyses in rat brains exposed to radiation and a second inflammatory insult, we revealed that radiation leads to molecular changes associated with microglial priming and validated this in normal appearing brain tissue of GBM patients treated with radiotherapy.

Importantly, we showed that radiation-associated microglial priming is a long-lasting response, which is in line with clinical observations in brain tumour patients of irreversible and progressive neurocognitive sequelae after treatment with radiotherapy. Recently, it has been demonstrated that the molecular state of microglia can be determined by microenvironmental changes^15^. Studies with DNA repair deficient *Ercc1* knock-out mice showed that generic and neuron-specific deletion of *Ercc1* induced microglial priming, whereas astrocyte- or microglia-specific deletion of *Ercc1* did not^20,26^. Neurons might be more sensitive to impaired DNA repair because they preferentially express a relatively large percentage of long genes, which generally acquire more DNA lesions than shorter genes, compared to other non-neuronal cell types^27,28^. These data suggest that the cause of persistent microglial priming induced by genotoxic radiation damage could be linked to neuronal DNA damage rather than resulting from a direct effect on the microglia.

The extent of radiation-associated microglial priming might also be affected by the severity and frequency of the initial radiation stimulus. In this study, we irradiated rats with a single fraction of radiation, mostly of 14 Gy, which is comparable to a total clinical fractionated dose that brain tumour patients generally receive^29–31^. Radiation dose fractionation with multiple lower radiation doses could change the nature and extent of the microglial response. Subsequent radiation exposures could desensitise microglia and cause tolerance instead of priming, as a single LPS challenge has been found to dampen the microglial response to subsequent LPS treatments^12,13^. In this study, we found that exposure to multiple lower radiation doses still led to microglial priming.

Due to high survival rates ofmore than 70%, radiotherapy-induced side effects are of particular concern in paediatric brain tumour survivors^32^. Here, we observed that radiation seems to affect the expression of microglial priming genes to a lesser extent in juvenile rats compared to adult rats. An increase in proinflammatory microglia with increasing irradiation age in rodents has been previously reported^33,34^. Radiation-affected microglia might have a greater impact on neurogenesis and phagocytosis at a younger age, while leading to a more pronounced neuroinflammatory response at an older age^35^.

In agreement with the animal data, we observed that the genes associated with the radiation response, including microglial priming genes, were upregulated in normal appearing brain tissue of GBM patients compared to control, indicating that microglial priming could be induced by brain cancer treatment in humans. As these patients did not only receive fractionated radiotherapy, but also chemotherapy, we cannot link the increase in microglial priming genes solely to radiotherapy. However, despite the fact that we compared rat microglial gene expression to human bulk gene expression, the correlation between the two groups is remarkable.

The notion that radiotherapy can induce persistent microglial priming is important to understand the development of cancer treatment-related neurotoxicities, in particular imaging changes, including pseudoprogression, and worsening of pre-existing neurological symptoms. Importantly, patients undergo other forms of immune-modulatory treatments in concomitance or after radiotherapy, including chemotherapy and immunotherapy, which could act as subsequent immune challenges and contribute to the worsening of neurotoxicities^36^. Notably, primed microglia have been hypothesised to be involved in delirium, a frequent neurocognitive complication experienced in patients with brain tumours^37,38^.

Together our findings indicate that radiotherapy treatment can shape the IIM of microglia leading to persistent priming. The knowledge that radiation makes microglia susceptible to secondary systemic immune challenges opens the way to the development of lifestyle changes and therapeutic strategies aimed at addressing systemic inflammation, and hence ameliorate radiotherapy-induced neurotoxicity and its impact on patients’ quality of life.

## Materials and methods

### Animals

Male Wistar (Hsd/Cpb:WU) rats were conventionally housed in groups under environmentally controlled conditions (temperature of 21°C and humidity of 55%) with a 12 hour light/dark cycle (light 08:00-20:00 in winter and 07:00-19:00 in summer) and ad libitum chow and water. Animals were randomly assigned to experimental groups. The animal procedures were performed in the Central Animal Facility of the UMCG according to the guidelines from Directive 2010/63/EU of the European Parliament on the protection of animals used for scientific purposes. The experiments were approved by the CCD (license # AVD1050020184808) and the Animal Care and Use Committee of the University of Groningen.

### Human post-mortem brain tissue

The use of post-mortem brain samples from deceased individuals was approved by the NIH NeuroBioBank (provider institution) and the University Medical Center Groningen (UMCG; recipient institution). RNA-seq data and the same paraffinized human post-mortem brain samples of Ainslie et al.^25^ were used for this study. A description of the included patient samples can be found in Table S4.

### Irradiation

The whole brain of the rats (males, mean weight 296 g ± SD 21 was irradiated with 14 Gy X-rays (X-RAD 320, Precision X-ray). Some animals were also irradiated with 10 or 18 Gy for the dose response study. In addition, to compare the response of a fractionated dose to a single dose, rats were irradiated with 5.8 Gy daily for five days. This fractionated dose is equivalent to a single dose of 14 Gy as calculated by the biologically effective dose equation with an α/β ratio of 2^29^. For the irradiation procedure, rats were anaesthetised using isoflurane (5% induction, 1.5-2% maintenance) and hung from their teeth on a specialised holder allowing the precise irradiation of the whole brain through a dedicated collimator on the dorsal side of the animal’s head^39^. Juvenile male rats at postnatal day 28 (P28) were irradiated in the same way using a collimator specifically designed to irradiate the whole brain of rats at this age. The same procedure was followed in sham-irradiated controls without irradiating the animals.

### LPS immune challenge

For the LPS immune challenge experiment, rats were randomly divided into one of four groups (*n* = 4): sham-irradiated controls that received phosphate-buffered saline (PBS) (control + PBS) or lipopolysaccharide (LPS) (control + LPS) and irradiated animals that received PBS (radiation + PBS) or LPS (radiation + LPS). Six weeks after irradiation, rats were intraperitoneally (i.p.) injected with PBS or with 1 mg/kg LPS in PBS, returned to their home cage, and perfused and sacrificed after four hours.

### Sacrifice procedure

Rats used for RNA sequencing were sacrificed 4 hours after LPS or PBS injections by perfusion and termination under isoflurane anaesthesia (5% induction, 1.5-2% maintenance). The rats used for qPCR analysis or microscopy were sacrificed 48 hours, 6 weeks, 12 weeks or 50 weeks post irradiation. At the specified time points, these rats were perfused and sacrificed under dexmedetomidine-ketamine anaesthesia. Their brains were isolated and either stored in medium for subsequent microglia isolation, dissected and snap frozen, or fixed in 4% paraformaldehyde for 48 hours and embedded in paraffin.

### Microglia isolation

After termination, the brains were isolated and kept in medium A (HBSS (Gibco, cat#14170-088) with 0.6% glucose (Sigma, cat#G8769) and 7.5 mM HEPES (Lonza, cat#BE17-737E) on ice. Microglia were isolated as described previously^40^, with all steps performed on ice or at 4^°^C. In short, the brain tissue was homogenised with a Potter-Elvehjem tissue grinder, filtered through a 108 µM strainer and pelleted by 10 minutes of centrifugation at 300 RCF. The pellet was homogenised in a 24% Percoll gradient, covered with PBS and centrifuged at 950 RCF for 40 minutes (accelerate: 4, break: 0) to separate the cells from the myelin. The cell pellet was incubated with anti-CD16/CD32 (eBioscience, cat#14-0161-85, RRID:AB_467134, 1:100) for 10 minutes to block Fc receptors. Cells were then incubated with CD45-Alexa Fluor 647 (Bio-Rad Laboratories, cat#MCA43A647, RRID:AB_321197, 1:100) and CD11b-FITC (Bio-Rad Laboratories, cat#MCA619F, RRID:AB_321303, 1:100) for 30 minutes before washing the cells. In order to select for live cells, DAPI (4’,6-Diamidino-2-Phenylindole, 200 ug/ml, Dilactate, cat#42280) was added to the samples, and subsequently microglia were sorted by gating for DAPI^neg^ CD11b^high^ CD45^int^ cells using the Beckman Coulter MoFlo XDP fluorescence-activated cell sorter (Figure S1). For RNA sequencing, 200.000 microglia per sample were sorted in siliconized Eppendorf tubes (Sigma, cat#T3406-250EA) in RLT+ buffer (from RNeasy Plus Micro Kit (Qiagen, cat#74034)), then snap frozen and stored at -80 ^°^C before RNA isolation.

### RNA isolation and RNA sequencing

Microglial RNA was isolated using the RNeasy Plus Micro Kit (Qiagen, cat#74034) according to the manufacturer’s instructions. RNA concentrations were measured using Agilent ScreenTape System. The sequencing libraries were generated by Genomescan using the NEBNext Low Input RNA Library Prep Kit from Illumina (New England Biolabs, cat#E6420S/L). In short, cDNA was reverse transcribed from mRNA with an oligo(dT) primer and a template-switching oligo (TSO) and amplified by PCR with sequencing primers. The quality, concentration and molarity of the libraries were then measured with the Fragment Analyzer HS NGS Fragment Kits (Agilent, cat#DNF-474). All libraries were pooled equimolar and sequenced on NovaSeq6000 and NovaSeq control software NCS v1.7 and the Illumina data analysis pipeline RTA3.4.4 and Bcl2fastq v2.20 for the following image analysis, base calling, and quality check.

### RNA sequencing analyses

Quality control of the generated sequencing data was performed with FastQC (v0.11.9) and FastQA (v3.1.27) followed by adapter trimming and quality filtering with fastp (v0.23.2.)^41^ Alignment with the Ensembl genome *Rattus norvegicus* (mRatBN7.2) was performed with STAR2 (v2.7.10)^42^, and transcripts were counted with HTSeq (v2.0.2)^43^. The following RNA-seq analyses were performed in the R environment (v.4.1.1) as previously described^44^. After filtering genes with counts <5, normalisation, transformation, and differential gene expression analysis were performed with edgeR (v3.36.0)^45^ and limma (v3.50.3)^46^. The transposed form of log counts per million (logCPM) matrix was used for the principal component analysis (PCA). Differentially expressed genes (DEGs) analysis was performed with a threshold of log^2^ fold change (LFC) > 2 and adjusted p-value < 0.05. Gene ontology (GO) enrichment analysis and gene set enrichment analysis (GSEA) were performed by using clusterProfiler (v3.0.4)^47^.

### RNA isolation and qPCR

Brain tissue from the anterior cortex of the rats was sectioned into 40 µm thin sections, after which RNA was isolated using the RNeasy Lipid Tissue Mini kit (Qiagen, cat#74804) according to the manufacturer’s instructions. RNA was transcribed to cDNA with M-MLV reverse transcriptase (Invitrogen, cat#28025013) according to the manufacturer’s instructions. qPCR was performed using iQ SYBR Green Supermix (Bio-Rad, cat#170-8885) and run in triplicate on a Bio-Rad real-time PCR system. Relative mRNA expression was calculated with ΔΔCT method using *Ywhaz* as internal control. Primer sequences are listed in Table S3.

### Immunofluorescence staining of rat brain tissue

Five μm paraffin sections of rat brain samples were dewaxed, and antigen retrieval was performed by boiling the sections for 10 minutes in 10 mM Sodium citrate with 0.05% Tween. The sections were washed and blocked in PBS with 2% donkey serum and 2% bovine serum albumin for an hour and incubated with rabbit anti-IBA1 (WAKO, cat#019-19741, RRID:AB_839504, 1:500) and goat anti-GPNMB (R&D Systems, cat#AF2550, RRID:AB_416615, 3µg/ml) primary antibodies in PBS with 2% donkey serum overnight at 4°C. After washing, the sections were incubated for an hour at room temperature with the secondary antibodies Alexa Fluor 488 (Invitrogen, cat#A-21206, RRID:AB_2535792, donkey anti-rabbit) and Alexa Fluor 568 (Invitrogen, cat#A-11057, RRID:AB_2534104, donkey anti-goat) in PBS. Nuclear staining was performed using DAPI.

### Immunofluorescence staining of human brain tissue

Five μm paraffin sections of human brain samples were dewaxed, and antigen retrieval was performed by boiling the sections for 3.5 minutes in HistoVT One (Nacalai tesque, cat# 06380-05). The sections were incubated with Sudan Black (0.5% in 70% EtOH) for 5 minutes, before they were washed and blocked in PBS+ (PBS + 0.3% Triton) with 2% donkey serum and 2% bovine serum albumin for an hour. The rest of the staining procedure was performed as described above for the rat brain samples with the exception of the primary antibodies, which were diluted in PBS+ instead of PBS.

### Microscopy and image analysis

For quantification of the GPNMB immunofluorescence staining, imaging was performed using a Leica DM6B microscope at 40x magnification. Five images per animal of the same region of the frontal cortex were made for the quantification of the rat microglial GPNMB expression. For the quantification of human GPNMB expression in microglia, eight images of normal appearing brain tissue of glioblastoma (GBM) patients and matching controls were made per patient (Table S4). Microglia, positive for both IBA1 and DAPI, were counted manually, and of these microglia, the percentage of GPNMB positive microglia was calculated. Representative microscopic images (Figure 2, 4) of the same region were taken using the Leica SP8X DLS confocal microscope at 40x magnification.

### Statistical analysis

Statistical analyses were performed by GraphPad Prism 8 (GraphPad Software Inc.). For the comparison of the Z-score mean differences of multiple groups in RNA-seq, two-way analysis of variance (ANOVA) followed by Tukey’s multiple comparisons test was used. For the comparison of the mean differences of relative RNA expression of multiple groups in qPCR at different time points post irradiation, radiation doses, dose fractionation schemes and ages at the time of irradiation, two-way ANOVA followed by Sidak’s multiple comparisons test with log_2_ transformed data, one-way ANOVA followed by Tukey’s multiple comparisons test with log_2_ transformed data, one-way ANOVA followed by Sidak’s multiple comparisons test with log_2_ transformed data, and two-way ANOVA followed by Tukey’s multiple comparisons test with log_2_ transformed data were performed, respectively. F-statistics was used to test the significance of regression coefficients in linear regression models in the R environment (v.4.1.1). For the comparison of percentages of GPNMB-positive microglia, a two-tailed T-test was used. All of the statistical details of experiments can be found in the Figure legends. We defined that a p-value less than 0.05 is statistically significant.

## Acknowledgements

This work was supported by KWF Kankerbestrijding (project numbers 11148 and 12487 to L.B.), ZonMw Off Road and IBA (project number 451001001 to L.B.), SU2C-CRUK Pediatric Cancer New Discoveries Challenge Team Grant (project number SU2C#RT6186 to L.B.). Post-mortem brain tissues (frozen and fixed) were acquired from the NIH NeuroBiobank. The authors would like to thank Peter van Luijk for the collimator design, the flow cytometry unit at University Medical Center Groningen (UMCG) for FACS support and the central animal facility at the UMCG for animal support. Some figures were created using BioRender.com.

## Author contributions

Conceptualization, L.B., B.J.L.E, I.R.H., and S.M.K.; Methodology, D.C.V., T.O., L.B., B.J.L.E, I.R.H., S.M.K., and F.B.B.; Investigation, D.C.V., T.O., Y.J., G.P.L., L.R.N., F.B.B., A.C.S., N.B., J.S., U.B., and L.B.; Resources, A.P.A. and L.R.N.; Writing – Original Draft, D.C.V, T.O., and L.B.; Writing - Review & Editing, B.J.L.E, I.R.H., S.M.K., and R.C.P; Visualization, D.C.V, T.O., Y.J., and L.B.; Supervision, L.B., B.J.L.E, I.R.H., S.M.K., and R.C.P; Funding Acquisition, L.B.

## Declaration of interests

The authors declare no potential conflict of interest.

## Supplementary information

**Figure S1:**
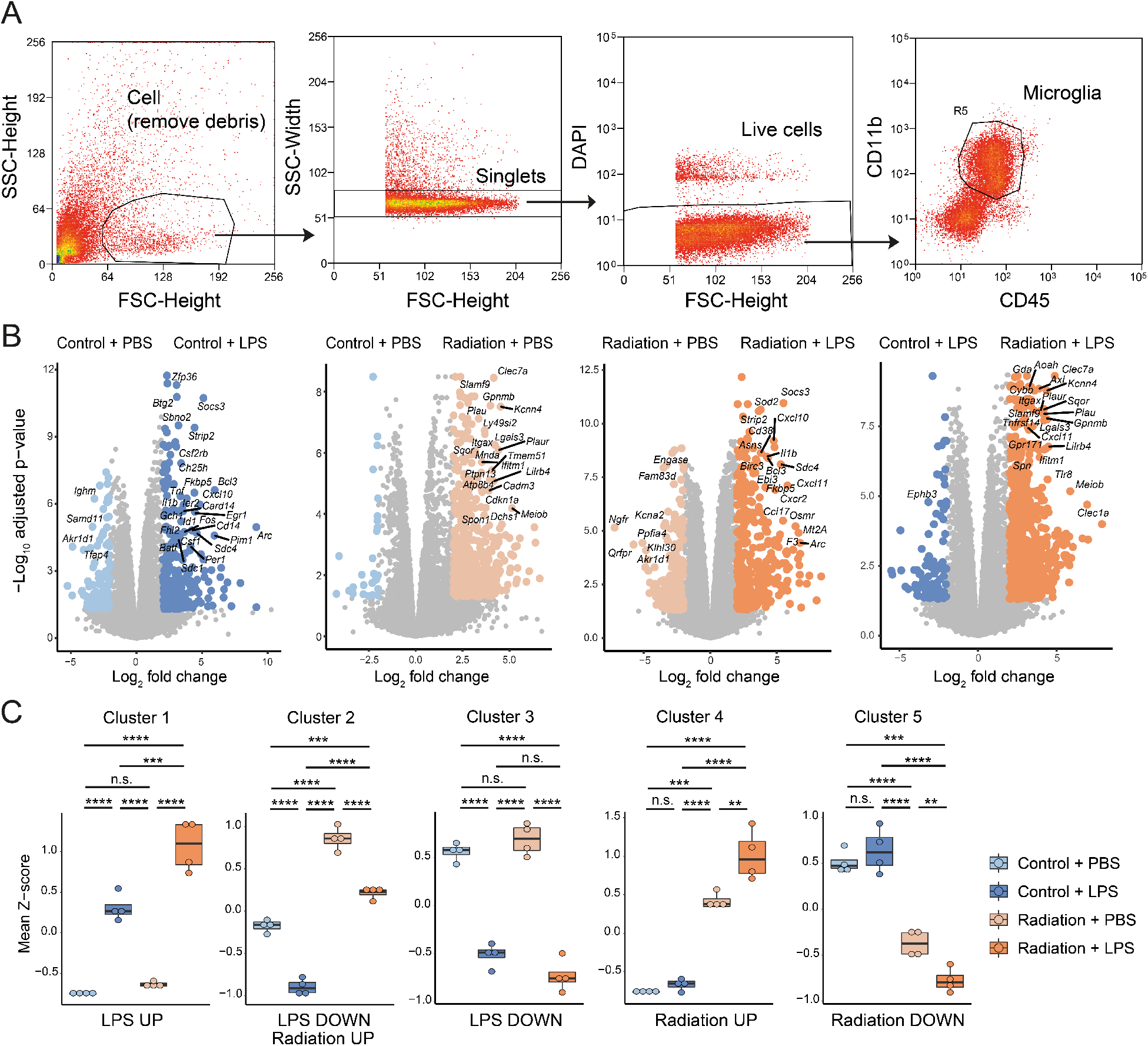
RNA-seq of microglia from irradiated rat brains exposed to an LPS immune challenge. (A) Representative FACS plots for the isolation of microglia (DAPI^neg^ CD11b^pos^ CD45^int^ cells). (B) Volcano plots showing differentially expressed genes (DEGs) (Log_2_ fold change (LFC) > 2, adjusted p-value < 0.05) between the conditions. (C) Box plots depicting the mean Z-score of genes in the clusters in each condition. Boxes are from the first quartile to the third quartile, and lines indicate the median. *n* = 4 animals per group. **** p <0.0001, *** p <0.001, ** p <0.01, n.s. not significant. Two-way ANOVA followed by Tukey’s multiple comparisons test was used for the comparison of the mean differences of multiple groups.

**Figure S2:**
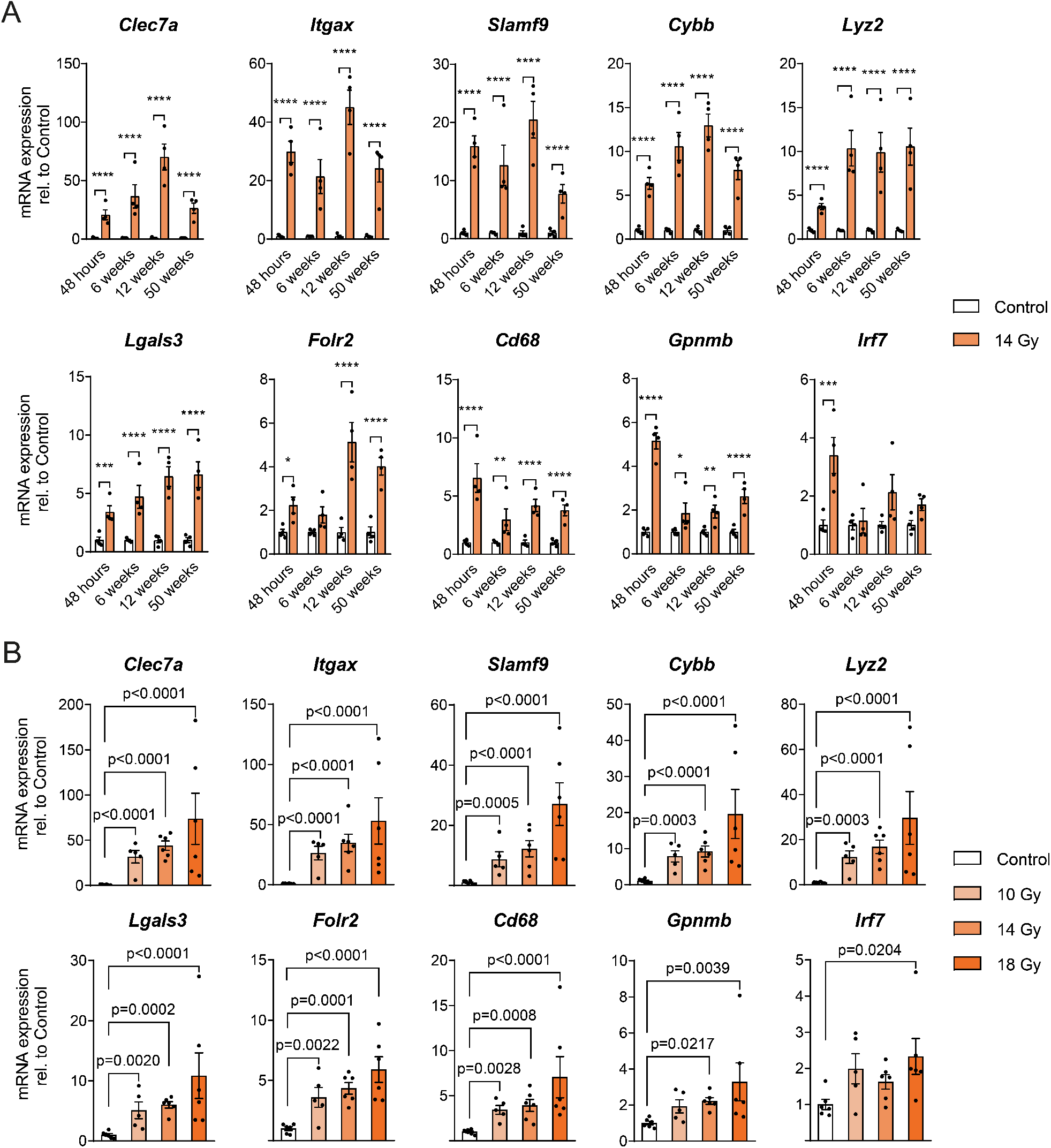
qPCR of microglial priming genes in irradiated rat brains at different time-points after irradiation and after different doses. Gene expression barplots of the microglial priming genes in irradiated or sham-irradiated rat brains (A) with series of time after irradiation (48 hours, 6 weeks, 12 weeks, 50 weeks) (B) or radiation dose (10 Gy, 14 Gy, 18 Gy). The y-axis shows the fold change of mRNA expression relative to corresponding controls. Bar graphs show mean ±SEM. **** p <0.0001, *** p <0.001, ** p <0.01, * p <0.05. (A) Two-way ANOVA followed by Sidak’s multiple comparisons test with log_2_ transformed data (*n* = 4 animals per group), (B) one-way ANOVA followed by Tukey’s multiple comparisons test with log_2_ transformed data (*n* = 5-6 animals per group) were performed.

**Figure S3:**
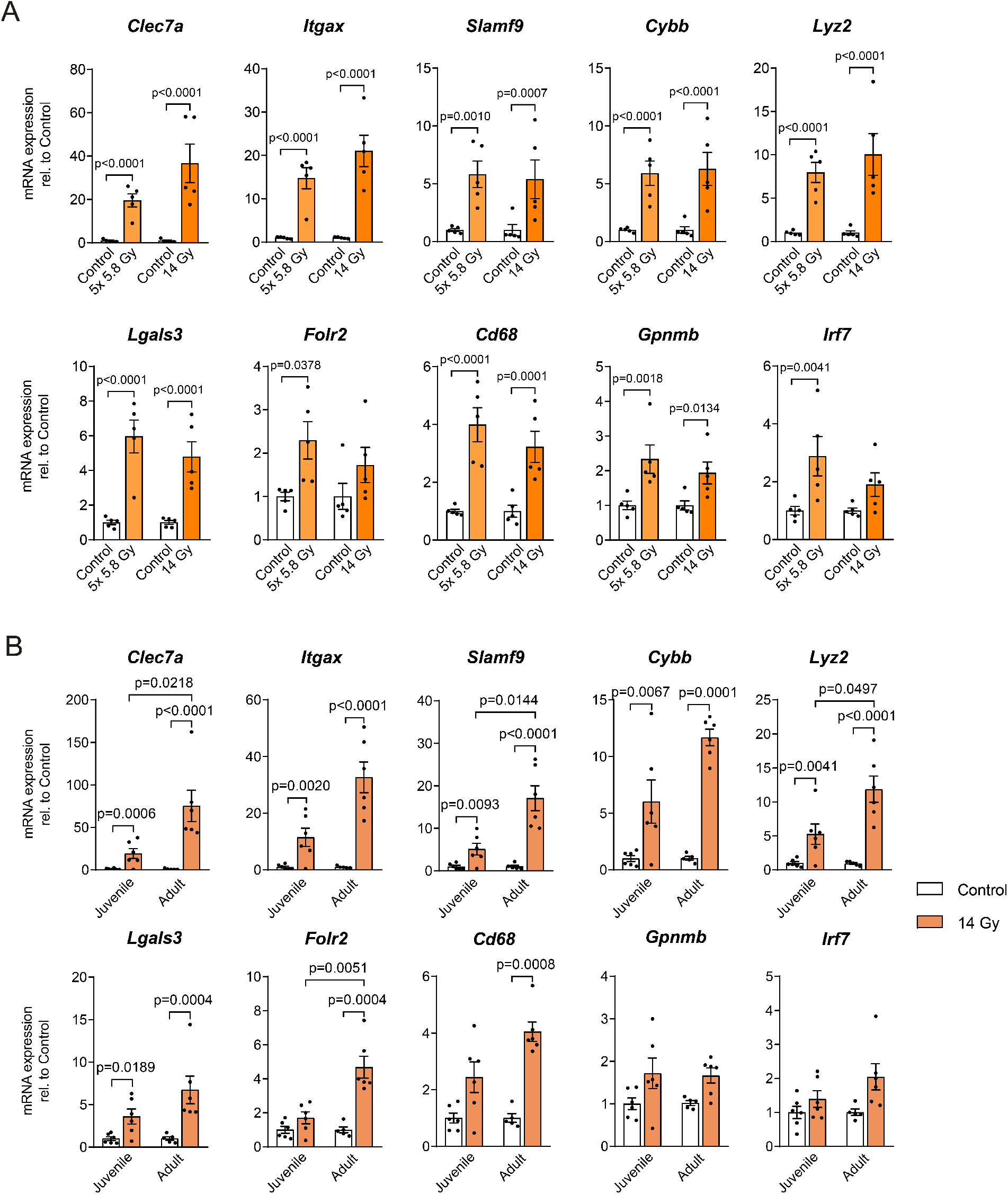
qPCR of microglial priming genes in irradiated rat brains after dose fractionation and different ages at the time of irradiation. Gene expression barplots of the microglial priming genes in irradiated or sham-irradiated rat brains (A) after irradiation with a fractionated dose of 5x 5.8 Gy or a single dose of 1x 14 Gy (B) or age at the time of irradiation (juvenile, old). The y-axis shows the fold change of mRNA expression relative to corresponding controls. Bar graphs show mean ±SEM. (A) One-way ANOVA followed by Sidak’s multiple comparisons test with log_2_ transformed data (n = 5 animals per group). The 5x 5.8 Gy group is normalised to the relative control group that received isoflurane anaesthesia 5 times. No significant differences were found between the 1x and 5x anaesthesia group (data not shown). (B) two-way ANOVA followed by Tukey’s multiple comparisons test with log_2_ transformed data (*n* = 5-6 animals per group) were performed.

**Figure S4:**
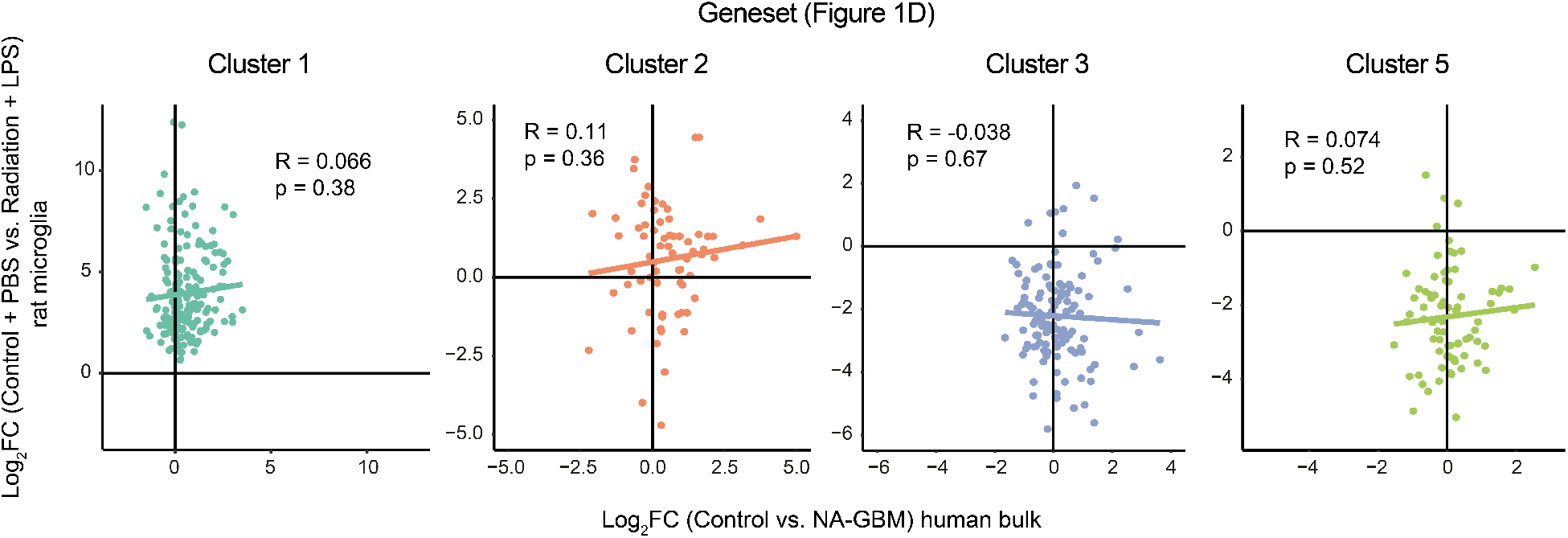
Gene cluster analysis in normal appearing brain tissue of GBM patients. Scatter plot depicting a correlation of gene expression enrichment of cluster 1, 2, 3, and 5 genes in figure 1d between the rat microglia-specific transcriptomic dataset (Control + PBS vs. Radiation + LPS) and the human bulk transcriptomic dataset (Control vs. NA-GBM) from Ainslie et al., 2022. F-statistics was used to test the significance of regression coefficients in linear regression models.

**Table S1: A list of the detected DEGs by the pairwise comparisons (related to figure 1)**

See excel file “Supplement Table S1 S2.xlsx”

**Table S2: A list of the DEGs in each cluster (related to figure 1)**

See excel file “Supplement Table S1 S2.xlsx”

**Table S3:**
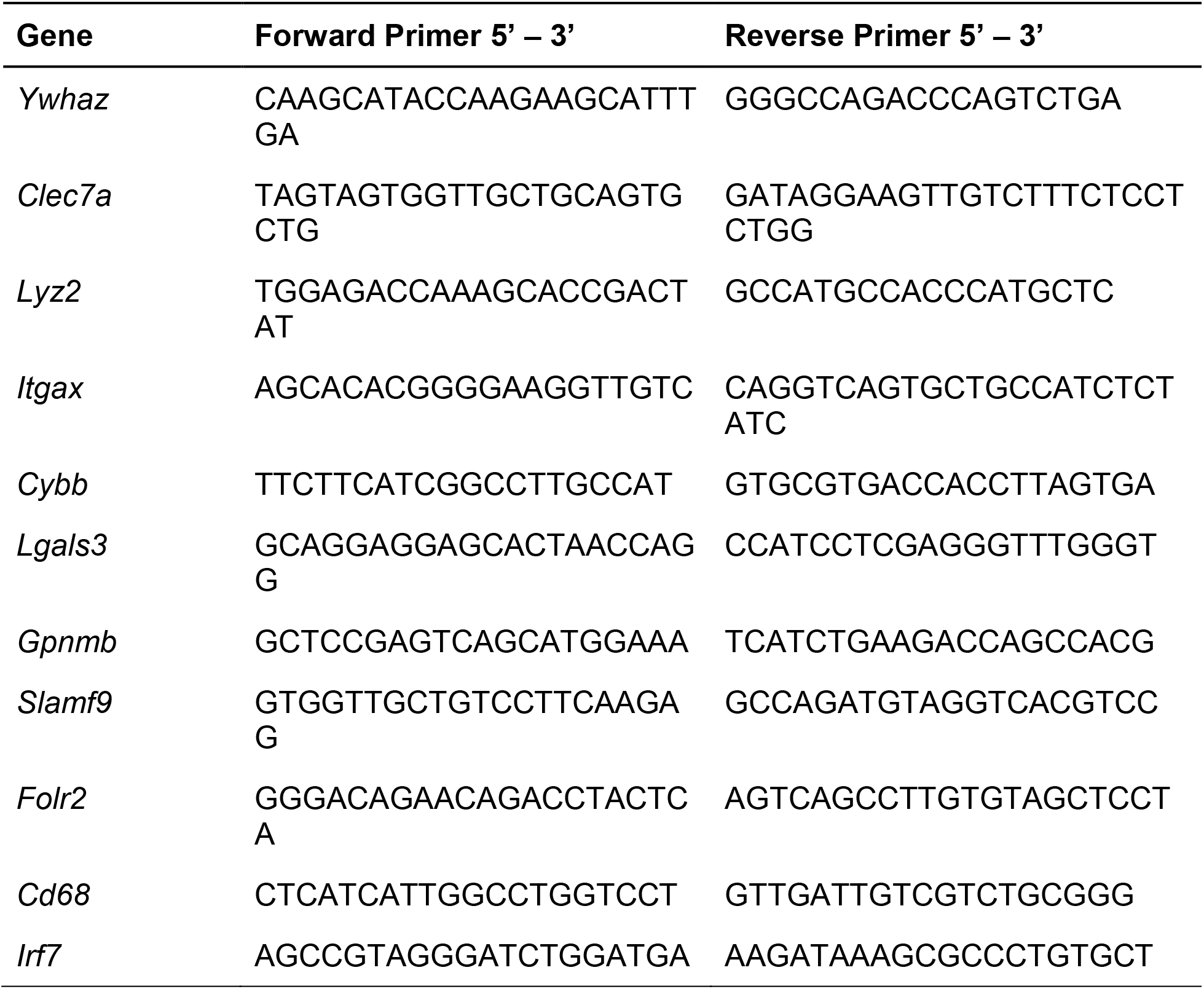
qPCR primer sequences.

**Table S4:** Description of included patient samples. Table describing the patient conditions, sample reference, age, sex assigned at birth, post-mortem interval (PMI), and whether samples were included in the RNA sequencing analysis (RNA-seq) or GPNMB staining (Staining).

| *Patient condition* | *Sample reference* | *Age* | *Sex* | *PMI (hours)* | *RNA-seq* | *Staining* |
| --- | --- | --- | --- | --- | --- | --- |
| *Unaffected control* | Control_1 | 54 | Female | 6 | Yes | Yes |
|  | Control_2 | 60 | Male | 37 | Yes | No |
|  | Control_3 | 54 | Male | 9 | Yes | No |
|  | Control_4 | 50 | Female | 15 | Yes | No |
|  | Control_5 | 49 | Female | 26 | Yes | Yes |
|  | Control_6 | 46 | Female | 18 | No | Yes |
| *Glioblastoma multiforme* | NA_GBM_1 | 62 | Male | 7 | Yes | No |
|  | NA_GBM_2 | 57 | Female | 8 | Yes | No |
|  | NA_GBM_3 | 55 | Female | 21 | Yes | No |
|  | NA_GBM_4 | 60 | Female | 21 | Yes | No |
|  | NA_GBM_5 | 49 | Female | 3 | Yes | Yes |
|  | NA_GBM_7 | 55 | Female | 18 | No | Yes |
|  | NA_GBM_9 | 47 | Female | 71 | No | Yes |

## Notes

### Competing Interest Statement

The authors have declared no competing interest.

